# Cytolethal Distending Toxin Enhances *Escherichia coli* Urinary Tract Infection

**DOI:** 10.64898/2026.02.09.704878

**Authors:** Santosh Paudel, Benjamin E. Curtis, Harry L. T. Mobley, Melanie M. Pearson, Mark T. Anderson

**Affiliations:** Department of Microbiology and Immunology, University of Michigan Medical School, Ann Arbor, Michigan, USA; ULAM Pathology Core, University of Michigan Medical School, Ann Arbor, Michigan, USA

## Abstract

Cytolethal distending toxin (CDT) is a virulence factor produced by several gram-negative bacteria, including *Escherichia coli*, the most prevalent etiological agent of urinary tract infections (UTI). CDT causes DNA damage in eukaryotic cells, impairing host defenses by disrupting epithelial barriers, suppressing acquired immunity, and promoting pro-inflammatory responses. *E. coli* strains encoding CDT have been previously identified in samples from UTI patients, yet the specific function of CDT in the development or progression of UTI remains undefined. In this study, we used a mouse model of ascending UTI to determine the role of CDT during infection. An *E. coli* mutant strain lacking the *cdtABC* locus was generated and combined with wild-type bacteria to co-infect mice via transurethral inoculation. At 1-day post-inoculation, competitive indices demonstrated a significant disadvantage for the *cdt* mutant in urine, bladder, and kidneys. Single strain infections were also performed as a further assessment of CDT impact, demonstrating that the *cdt* mutant had reduced kidney colonization indicative of CDT contributions to ascending infection. Histopathological analysis of the urinary bladder and kidney tissues from mice infected with CDT-encoding *E. coli* demonstrated higher levels of inflammation and tissue damage within the kidneys at both 1 and 7 days post-inoculation and in the bladder at 7 days post-inoculation compared to mice infected with *cdt* mutant bacteria. Collectively, these findings identify a critical contribution of CDT to the progression of UTI.

## Introduction

Cytolethal distending toxin (CDT) is a bacterial genotoxin secreted by numerous pathogenic Gram-negative bacteria including *Aggregatibacter actinomycetemcomitans* (1), *Campylobacter jejuni* (2), *Escherichia coli* (3), *Haemophilus ducreyi* (4), and *Salmonella spp* (5, 6). In *E. coli*, five variants of CDT (CDT-I through CDT-V) have been identified and differentiated by their genomic location (7). CDT-II is chromosomally encoded (8), CDT-III is located on a pVir plasmid (9), CDT-I and CDT-IV are associated with lambdoid prophage (10), and CDT-V can be found within chromosomal genes flanked by bacteriophage P2 and lambda-like sequences or in inducible bacteriophages (11). All variants have been observed in pathogenic *E. coli* strains of both intestinal and extraintestinal origin (3, 12, 13).

CDT is a heterotrimeric protein complex composed of the subunits CdtA, CdtB, and CdtC. The CdtA and CdtC subunits facilitate host cell binding and cellular internalization of the CdtB subunit via interaction with cholesterol rich membrane domains (14), and in some species, through binding of CDT-containing extracellular vesicles to host cell glycans (15). CdtB is the catalytically active subunit of the toxin, possessing DNase (16) and phosphatase activity (17). CdtB DNase activity induces double-strand DNA breaks in mammalian cells, triggering host DNA damage response pathways and leading to cell cycle arrest, apoptosis, and inflammation (16). The phosphatase activity is less well characterized and is thought to support toxin entry and nuclear trafficking (17, 18).

*E. coli* CDT induces characteristic cytopathic effects across several epithelial cell types (*e*.*g*. HeLa, U2OS, RKO, Hep-2, HCECs), including megalocytosis, chromatin fragmentation, and multinucleation (3, 19, 20). These phenotypes stem largely from G2/M cell cycle arrest and defective DNA repair, resulting in genomic instability and pro-tumorigenic signaling (21, 22). Although CDT-mediated cytotoxicity has been extensively described in intestinal and colonic epithelial models (19, 23, 24), including CDT-positive *E. coli* isolated from the gastrointestinal tract (3, 19), the contribution of CDT to urinary tract infection (UTI) remains undefined.

UTIs represent a significant global health burden which has become increasingly prevalent over the past 30 years (25). Uropathogenic *E. coli* (UPEC) accounts for about 75% of uncomplicated and 65% of the complicated UTI cases (26). UPEC employ a diverse array of virulence factors that contribute to adherence, colonization, immune evasion, and tissue damage within the urinary tract (27–30). CDT producing *E. coli* has been identified among UTI isolates at a rate of approximately 10% in limited sample sizes (13, 31, 32), but its role in the urinary tract remains undefined. This study investigates the contribution of CDT to UPEC virulence during UTI by assessing bacterial burden, inflammatory responses, and tissue pathology in the bladder and kidneys. Understanding the contribution of CDT to the pathogenesis of the urinary tract will advance our understanding of the mechanisms that drive UTI development and progression.

## Results

### Identification of UPEC *cdt* genes

We identified putative *cdt* genes via homology search from the previously reported genome sequences of two uncomplicated *E. coli* UTI isolates, HM56 and HM57 (31) (Table 1). CDT subtype was inferred by aligning published CDT-typing primer sequences to the HM56 and HM57 genome assemblies. The CDT-IV specific primers matched the *cdt* operon of HM56, while the CDT-I specific primers matched the *cdt* operon of HM57, consistent with classification as CDT-IV and CDT-I, respectively (13). Both CDT-I and CDT-IV have been reported to be carried within lambdoid prophage regions in the *E. coli* genome (10). The PHASTEST platform (33) was used to confirm prophage locations and annotate each of the intact prophage regions in HM56 (n=4 regions) and HM57 (n=4 regions). The *cdt* genes were encoded in region 4 for HM56 and region 3 for HM57 (Fig. S1).

**Table 1:** Bacterial strains and plasmids used in this study.

| Strain | Genotype | Source |
| --- | --- | --- |
| Bacteria |  |  |
| *E. coli* HM56 | Wild-type | {31633483} |
| *E. coli* HM56 | $\Delta cdt::kan^R$ | This study |
| *E. coli* HM57 | Wild-type | {31633483} |
| *E. coli* HM57 | $\Delta cdt::kan^R$ | This study |
| Plasmid |  |  |
| pKD4 | Source of FRT-*kan*<sup>R</sup>-FRT marker | {10829079} |
| pSIM18 | Encodes $\lambda$ red recombineering system,<br>hygromycin<sup>R</sup> | {16750601} |
| pBBR1MCS-4 | Cloning and shuttle plasmid vector | { 8529885} |

To determine whether *cdt* genes are expressed under conditions that approximate the urinary tract environment, we quantitated the expression of *cdtB* in standard Lysogeny broth (LB) culture and human urine (hU). The relative cdtB expression in hU compared to LB was 1.76±0.10 and 1.35±0.11 (mean±SD) fold higher at times 1 and 4h, respectively (Fig. S2), indicating that the *cdtB* gene may be actively expressed during infection of the urinary tract.

### CDT confers a fitness advantage in the urinary tract

To initially assess the fitness contribution of CDT during UTI, we performed co-challenge infections with wild-type HM56 and a Δ*cdt* mutant (1:1) in a murine model. Urine, bladder, and kidneys were recovered from infected mice followed by the quantitation of wild-type and mutant bacteria at each site (Fig. 1A). The competitive index (CI) indicates a significant fitness disadvantage for Δ*cdt* mutant bacteria throughout the urinary tract at 24h post-inoculation (Fig. 1B). Relative fitness of the mutant was lowest in the bladder, with an 8-fold difference compared to wild-type bacteria.

**Fig. 1.**
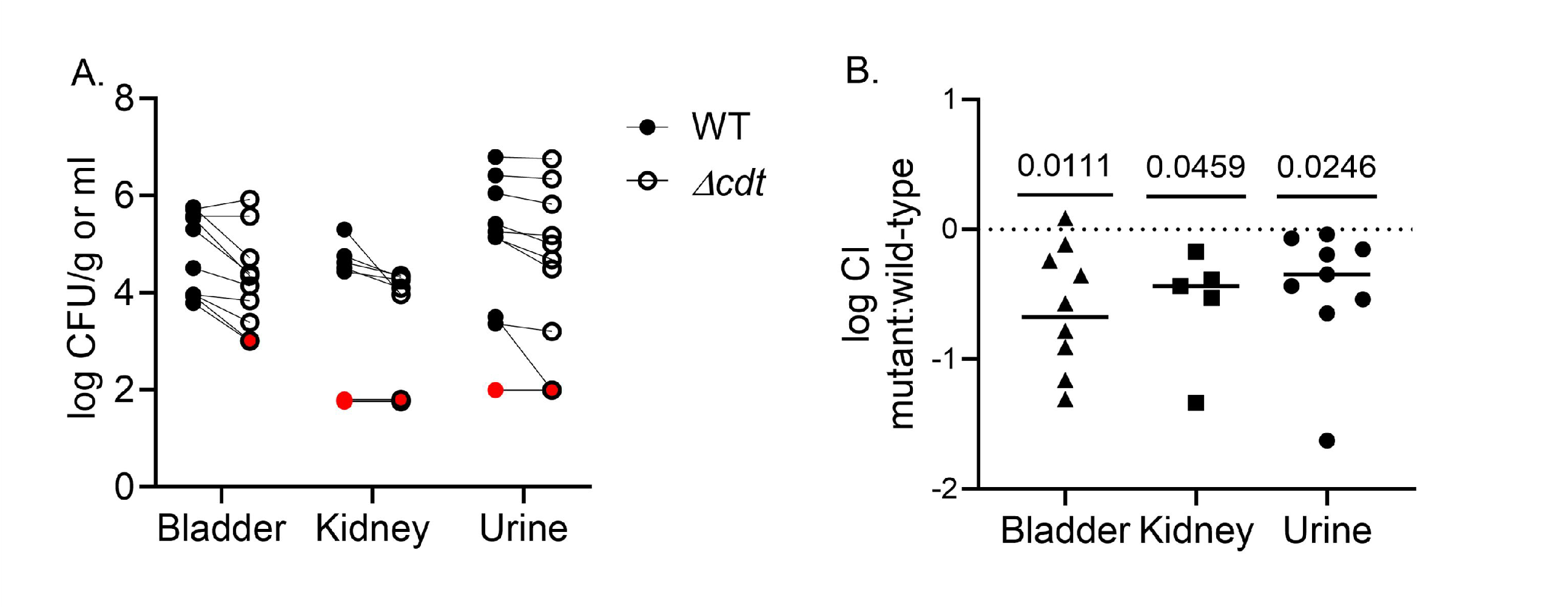
CDT contributes to UTI fitness of *E. coli*. CBA/J mice (n=10) were transuretherally inoculated with equal numbers of wild-type (WT) HM56 and Δ*cdt* mutant HM56. Urine, bladders, and kidneys were collected for CFU enumeration at 24 h post-inoculation. A. CFU of each strain recovered from individual mice. Red symbols indicate no CFU recovered after plating. B. CI of the Δ*cdt* mutant compared to wild-type. Statistical significance was assessed by a one-sample t-test, comparing the mean log CI (lines) to a hypothetical value of zero for neutral fitness (dotted line): Statistically significant if P-values < 0.05.

To determine whether the UTI fitness defect of the Δ*cdt* mutant was attributable to differences in bacterial growth rate, we compared the growth kinetics of the HM56 wild-type and Δ*cdt* mutant strains under *in vitro* and *in vivo* conditions. No significant difference in the population doubling time was observed between these strains in LB medium calculated from the CFU measured during the exponential growth phase from 1-3h post-inoculation, calculated at 25.5±1.2 and 25.7±3.4 minutes respectively (Fig. 2A). Similar experiments were also performed in human urine and RPMI medium, with minimal differences observed between the two strains in these conditions (Fig. S3). In preparation for measuring replication *in situ*, we also determined growth rates of bacteria in LB culture by our previously reported O:T^PCR^ method (34). O:T^PCR^ measurements, a proxy for the relative abundance of genome *origin* and *terminus* copies in the population, were consistent across both strains and showed similar replication dynamics over 6 h (Fig. 2A). As expected, doubling times calculated from the highest O:T^PCR^ values (at T=1.5 h) were also consistent at 19.7±1.9 and 18.1±0.3 minutes for wild-type and the Δ*cdt* mutant strains, respectively. Using the O:T^PCR^ method, we then investigated the growth rates of bacterial populations recovered from the urine of infected mice and no significant difference was found in the doubling times of wild-type and the Δ*cdt* mutant was detected at 6 or 24 h (Fig. 2B). Growth was fastest for both strains 6 h after inoculation (39.3 and 38.1 min doubling time) and slowed by 24 h (45.6 and 50.9 min doubling time) (Fig. 2B). Together, these results demonstrate that there was no inherent growth limitation of the Δ*cdt* mutant under the conditions tested and that the fitness defect of this strain is not attributable to differences in bacterial replication.

**Fig. 2:**
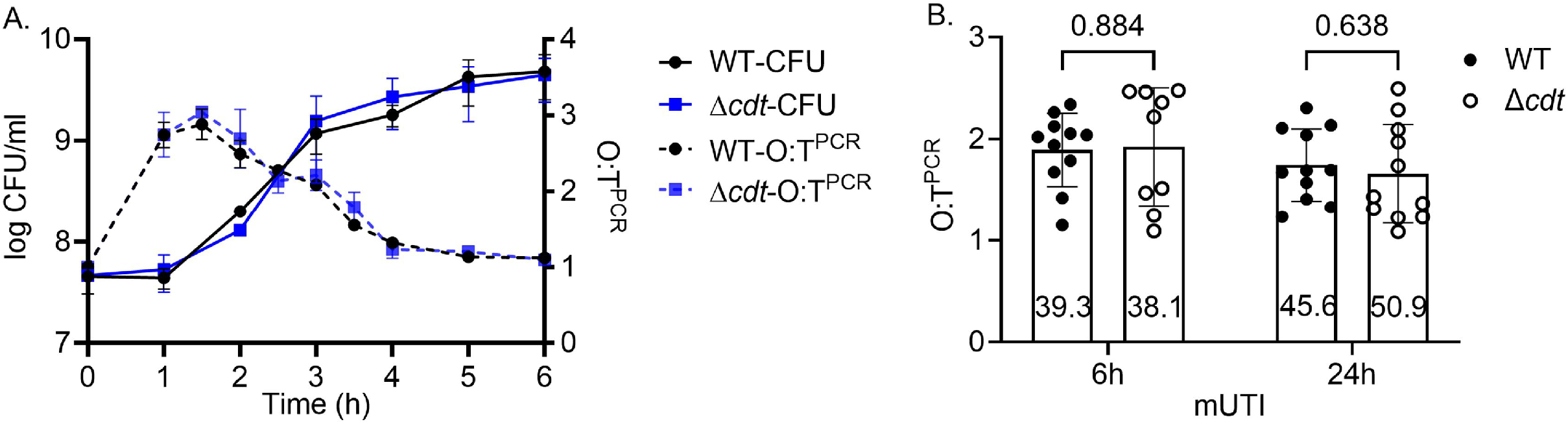
Wild-type HM56 and the Δ*cdt* mutant have similar growth kinetics in culture and during infection. A. CFU from wild-type (WT) and Δ*cdt* cultured in LB are plotted as the mean (n=2) ± the standard deviation or as O:T^PCR^ (n=9-11). B. O:T^PCR^ values from the urine of infected mice collected at 6 or 24 h is plotted with inset numbers indicating the doubling time derived from the median O:T^PCR^ (bars). Statistical significance was assessed using the Mann–Whitney U test. P-values are displayed on the graph, and neither was below the 0.05 significance threshold.

### HM56 CDT contributes to kidney colonization

We further examined the role of CDT in colonization of the urinary tract using a single-strain infection model of ascending UTI, tracking bacterial abundance in the urine, bladder, and kidneys over the course of a week (Fig. 3A). Both wild-type and Δ*cdt* mutant bacteria persisted in the urine of mice throughout the seven-day infection period. However, urine bacterial burdens were highly variable and there was no significant difference in CFU levels between the two strains at most time points (Fig. 3B). The one exception was observed at 3 days post-inoculation, where wild-type bacteria exhibited an approximately two-log higher median CFU compared to Δ*cdt* bacteria (Fig. 3B). Both strains individually were similarly able to colonize the bladder, and no significant difference was observed between wild-type HM56 and Δ*cdt* mutant CFU at one or seven days post-inoculation (Fig. 3C). In the kidney, wild-type bacteria exhibited a >3-log higher CFU burden compared to the Δ*cdt* mutant at one day after inoculation, representing a significant increase in either colonization or bacterial survival at this site (Fig. 3D). A trend toward decreased Δ*cdt* kidney CFU was also observed after 7 days; however, this difference was not significant. Overall, bacteria were recovered from a higher proportion of kidneys from mice inoculated with wild-type bacteria (1 day, 12/13; 7 days, 11/13) than from those inoculated with the Δ*cdt* mutant (1 day, 6/13; 7 days, 9/13). Together, these results demonstrate that CDT enhances kidney colonization by *E. coli*.

**Fig. 3.**
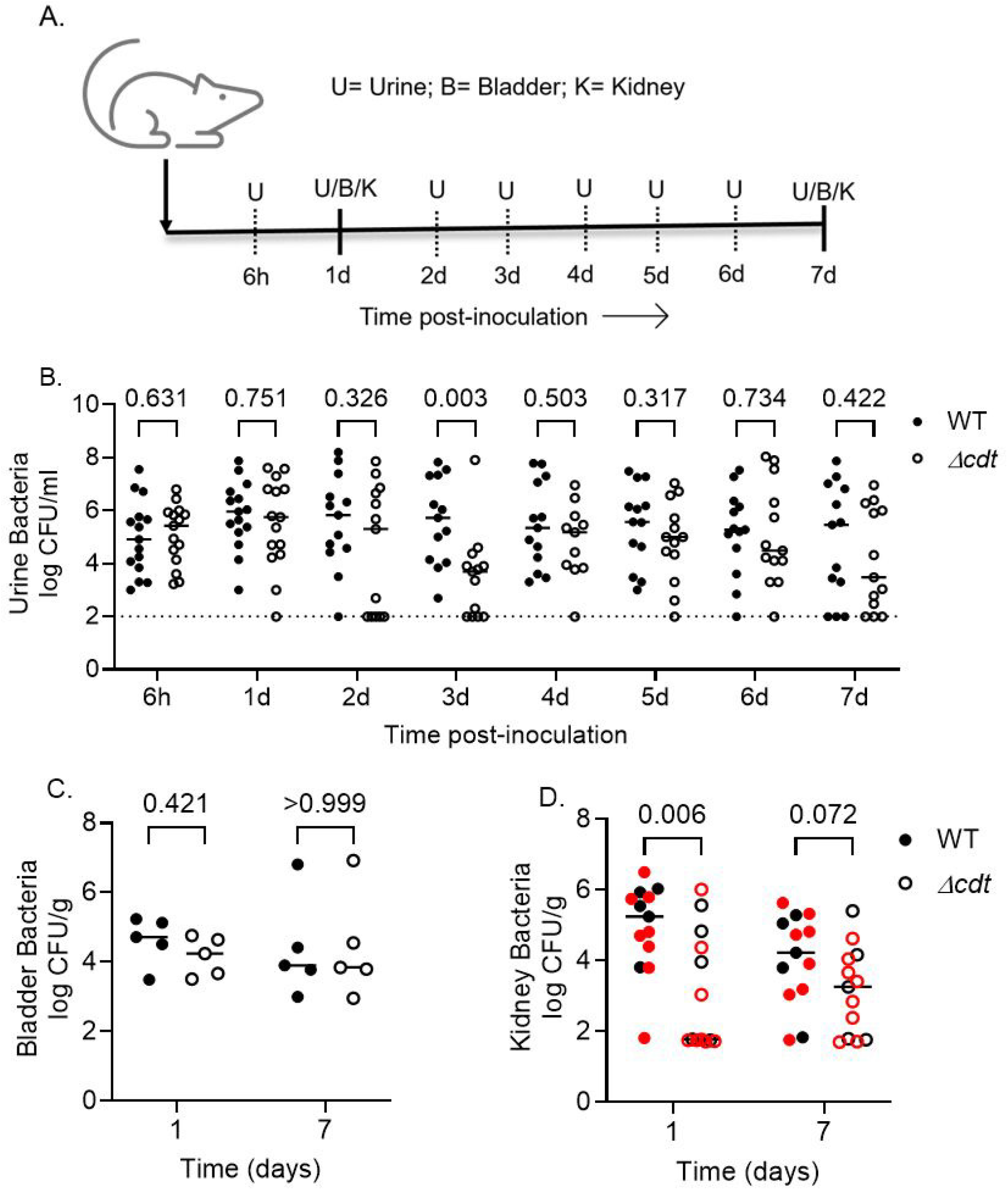
CDT contributes to short- and long-term kidney colonization in a single-strain UTI model. A. Timeline of infection and bacterial enumeration in female CBA/J mice transurethrally inoculated with wild-type (WT) HM56 and Δ*cdt* mutant strains. B-D. Bacterial burdens from individual mice in urine (B), bladder (C), and kidneys (D) (*n* = 5-15) with the median indicated by solid lines and the limit of detection (urine) indicated by the dotted line. The red symbols in panel D designate mice in which bacterial counts were derived from only one kidney; the remaining kidney was preserved for histology. Statistical significance was assessed using the Mann–Whitney U test. P-values are displayed in the graph, and values < 0.05 were considered statistically significant.

To determine whether CDT contributed to UTI in a second strain, HM57, a Δ*cdt* mutant derivative was constructed and used to infect mice. Unexpectedly, no significant difference in bacterial colonization was observed in the urine, bladders, or kidneys of infected mice up to 24h after inoculation (Fig. S4). The observed discrepancy in results between HM56 and HM57 suggests that the contribution of CDT to UTI may be influenced by the type of CDT variant encoded or dependent on the larger genetic differences between these two UPEC isolates that could affect pathogenesis. Notably, HM57 encodes at least two additional toxins, hemolysin and cytotoxic necrotizing factor, which are not present in HM56 genome (Table 2).

**Table 2:** Comparison of predicted toxins encoded in UPEC strains HM56 and HM57.

| Toxins | HM56 | HM57 |
| --- | --- | --- |
| Cytolethal distending toxin (*cdt*) | + | + |
| Hemolysin (*hly*) | - | + |
| Cytotoxic necrotizing factor (*cnf*) | - | + |
| Serine protease autotransporter (*vat*) | + | + |

+ Present; - Absent

### CDT induces inflammation in the bladder and kidney

As a genotoxic effector, CDT has the potential to impact the development of pathology in the urinary bladder and kidneys during UTI. Histological analysis revealed that mice infected with wild-type HM56 developed more severe inflammation and more extensive tissue changes in their kidneys and bladder compared to mice infected with the Δ*cdt* mutant strain (Fig. 4). Specifically, bladders of mice infected with wild-type bacteria displayed more pronounced mucosal hyperplasia with submucosal expansion by edema and infiltrates of inflammatory cells (Fig. 4A). Mucosal injury was also more common and characterized by vacuolated epithelial cells, frequent apoptotic bodies, and sloughed epithelial cells. Inflammation index scoring (Table S2, Fig. S5) demonstrated that 6/8 mice exposed to wild-type HM56 and 4/8 mice exposed to the Δ*cdt* strain showed some level of cystitis at one day post-inoculation, with a median severity score of 1.0 and 0.5, respectively (Fig. 4B). At 7 days post-inoculation the incidence of cystitis was significantly lower in mice infected with the Δ*cdt* mutant (2/8 mice, median severity = 0) compared to mice that received wild-type bacteria (8/8 mice, median severity = 1.0) (Fig. 4B). Importantly, the bacterial burden of both strains was similar in the bladder after 7 days (Fig. 3C), indicating that the differences in lesion severity between groups were unlikely due to differences in the abundance of bacteria at this site. CDT, therefore, contributes to HM56-induced pathology of the urinary bladder.

**Fig. 4.**
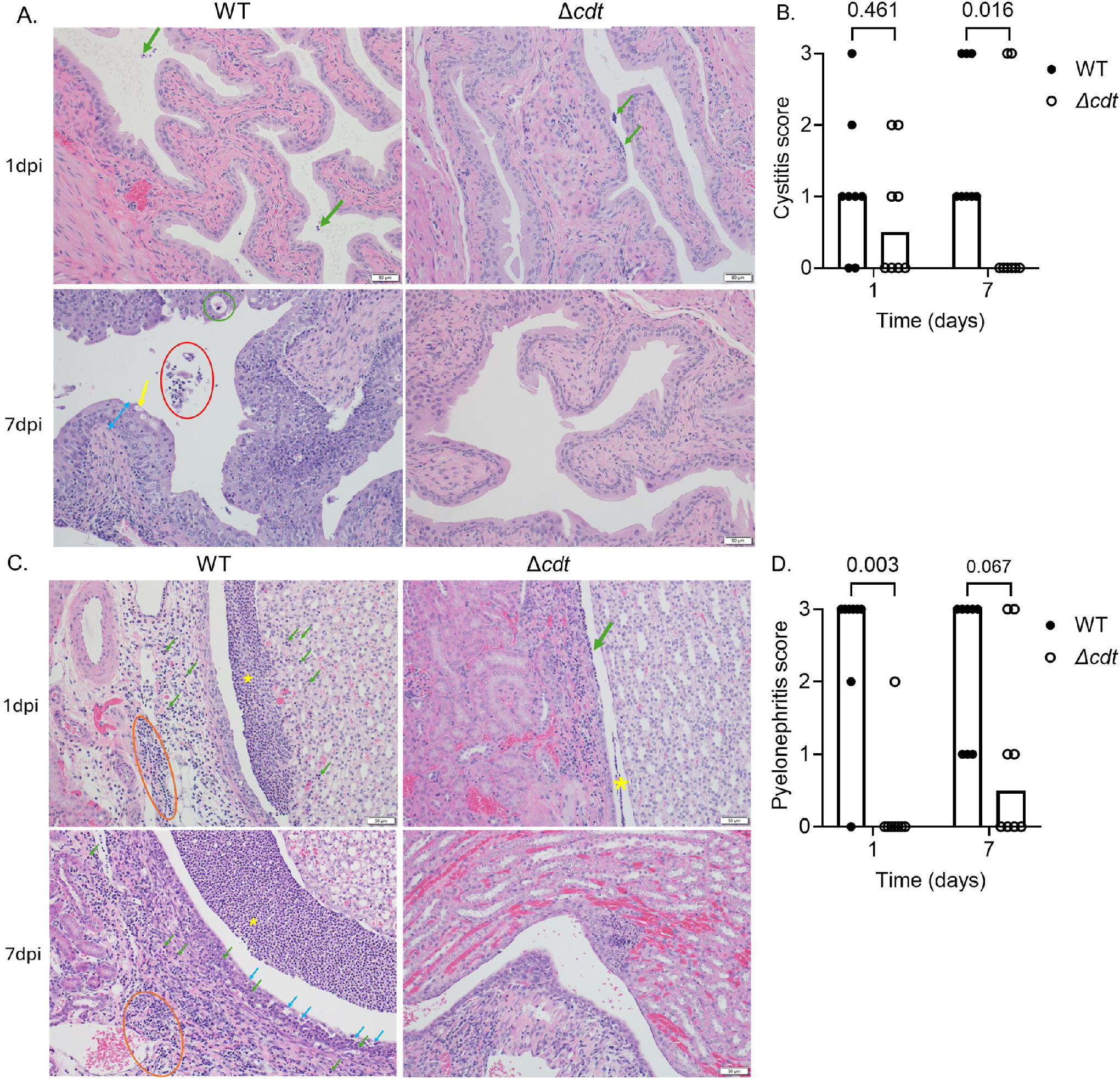
HM56 CDT contributes to cystitis and pyelonephritis pathology during UTI. A. Sections of urinary bladder from mice infected with wild-type (WT) or Δ*cdt* mutant bacteria were stained with hematoxylin & eosin to visualize tissue damage and inflammatory cells. B. The histology scores of individual bladder samples are presented with bars representing the median score. C. Kidney sections from infected mice were stained as described for bladders. D. Histology scores of individual kidneys with bars representing median scores. Images were taken at 200X magnification. Scale bar = 50 µm. Statistical significance in panels B and D was assessed using the Mann–Whitney U test. P-values are displayed on the graph, and values < 0.05 were considered statistically significant. Histology Key: Green arrow (→), PMN infiltrates; orange circle (**O**), lymphoplasmacytic infiltrates/cuffs; yellow asterisk (**\***), PMN rafts; blue arrow (→), urothelial (mucosal) injury; yellow arrow (→), vacuolation; green circle (**O**), apoptotic body; red circle (**O**), sloughed cells and bacteria; blue double arrow (↔), mucosal hyperplasia.

Mice exposed to wild-type HM56 also more frequently developed pyelonephritis, or ascending inflammatory lesions in the kidneys centered on the renal pelvis. Pyelonephritis lesions were characterized by prominent rafts of polymorphonuclear cells (PMN), injury of the urothelium with infiltration of PMNs into the renal parenchyma, and expansion of perivascular tissue by edema, PMNs, and lymphoplasmacytic cuffs (Fig. 4C). The inflammation index scoring (Table S2, Fig. S5) showed significantly lower levels of inflammation in the kidneys of mice infected with the Δ*cdt* mutant compared to wild-type at 1 day post-inoculation, and a trend toward lower inflammation at 7 days. Pyelonephritis was observed in 3/8 kidneys after one day (median severity, 0) and 2/8 kidneys (median severity, 0.5) after 7 days in mice inoculated with the Δ*cdt* mutant, compared to 7/8 (median severity, 3.0)) and 8/8 (median severity, 3.0) at one and seven days, respectively, for kidneys from mice infected with wild-type bacteria (Fig. 4D). These results suggest that CDT contributes to the development and severity of pyelonephritis during UPEC infection.

Genotoxic CDT activity may result in cell death via apoptosis (35). To preliminary assess this activity in the context of UTI, apoptosis was quantitated by immunohistochemical detection of active caspase-3 (casp3) (Table S3). Casp3 reactivity was not detected in bladders from mice infected with either wild-type HM56 or the *Δcdt* mutant (Fig. S6 A, B). Casp3 reactivity was detected within kidney sections and differed significantly between infection groups. Mice infected with the wild-type HM56 strain showed higher casp3 reactivity (median value of 2.0), with 5 of 6 mice positive, compared to mice infected with the *Δcdt* strain (median value of 0), in which only 1 of 6 mice was positive (Fig. S6 A, C). However, it should be noted that reactivity was largely restricted to degenerate PMNs within the rafts in the renal pelvis. So, it is not surprising that samples with the greatest severity of pyelonephritis also had the greatest number of casp3 reactive cells. Occasional individual tubular epithelial cells displayed moderate casp3 staining; however, diffuse or regionally extensive tubular involvement was not observed. Together, this suggests that the observed caspase-3 reactivity is more associated with the progression of inflammatory responses than with the early stages of infection.

## Discussion

CDT is a virulence factor that has been identified in multiple bacterial pathogens and associated with diverse infection conditions. It has been implicated in the pathogenesis of gastrointestinal disorders including colitis (36) and colorectal cancer (2). For *E. coli*, CDT was initially characterized in gastrointestinal isolates, but it has also been detected in strains recovered from patients with UTI (13, 31). A previous regional study reported that 8% of *E. coli* isolates from UTI cases (n=190) harbor CDT-encoding genes (13). In a previous study from our group, 14% of *E. coli* isolates obtained from uncomplicated UTI cases (n=14) were positive for CDT genes. Both strains from this study were recovered from patients with history of recurrent urinary tract infection (31). A more recent survey of UPEC virulence factors found the *cdtB* gene in 3% of UTI isolates (n=31) from women ≥65 years old compared to 0% (n=15) from asymptomatic bacteriuria cases (32). These data, together with our findings here, support the role of CDT as an accessory virulence factor that may contribute to the UTI in a subset of UPEC strains. However, broader functional characterization of UPEC CDT variants is warranted given the observed differences between HM56 (CDT-IV) and HM57 (CDT-I).

The results from our infection model demonstrate the importance of CDT in the fitness and colonization of UPEC during UTI. In the competitive infections, attenuation of fitness for the Δ*cdt* mutant in the urine, bladder, and kidneys provides compelling evidence that CDT provides a significant advantage for bacterial colonization or survival throughout the urinary tract. In our single-strain infections, bladder and urine UPEC burdens were comparable between wild-type and Δ*cdt* strains, but there was significantly reduced recovery of Δ*cdt* bacteria from the kidneys. Therefore, CDT may either facilitate dissemination to or survival in the upper urinary tract, potentially by modulating the host response or overcoming anatomical barriers. The basis for the fitness defect observed in urine and bladder during co-challenge but not in single-strain infections, despite comparable CFU levels in each, is currently unknown. However, these differences may result from the enhanced sensitivity of the co-challenge measurements or other unknown factors such as nutrient competition, antagonism, or differences in immune evasion.

Examination of infected bladder and kidney tissues showed that HM56 encoding *cdt* genes significantly contribute to urinary tract pathology by enhancing epithelial injury and inflammation. Previous studies have shown that CDT induces DNA damage leading to cell cycle arrest and apoptosis in epithelial cells, disrupting barrier integrity and facilitating inflammation (23, 37, 38). The damage of epithelial cells promotes robust lymphoplasmacytic infiltration and PMN accumulation, as observed here in kidneys and bladders infected with wild-type HM56 infection. These phenotypes were reduced in Δ*cdt* strain, consistent with the reports that CDT triggers pro-inflammatory responses and suppresses immune resolution (22, 24, 39, 40). The increased epithelial changes (mucosal hyperplasia, vacuolation, apoptosis, sloughing) and immune cell infiltration seen here also resemble hallmark pathological features linked to *Helicobacter hepaticus* CDT-mediated genotoxicity in the gastrointestinal tract (39).

Combined, our results demonstrate that CDT acts as a potent virulence factor in certain UPEC strains by facilitating bacterial colonization, playing a role in tissue damage and inflammatory pathology, and, together, exacerbating disease severity in the urinary tract.

## Materials and Methods

### Bacterial strains and mutant generation

*E. coli* strains HM56 and HM57, isolated from uncomplicated UTIs as reported elsewhere (31), were routinely cultured at 37℃ with aeration in lysogeny broth (LB) (41) at 200 rpm unless stated otherwise. The *cdtABC* locus was deleted (Δ*cdt*) using the bacteriophage lambda-red recombination system as previously described (42). Briefly, primers SP1/SP2 (HM56) and SP5/SP6 (HM57) (Table S1) containing 5′ 35-40 bp sequences homologous to the flanking regions of *cdt* locus were used to amplify the kanamycin resistance cassette of plasmid pKD4 (43). These PCR products were used to transform HM56 or HM57 harboring pSIM18 (44) by electroporation. Transformants were selected at 30℃ on LB agar plates supplemented with kanamycin (50 µg/ml), screened by endpoint PCR using primers SP3/SP4 (HM56) or SP8/SP9 (HM57) (Table S1), and confirmed by sanger sequencing (Eurofins). Prophage sequences in bacterial genomes were detected and annotated using PHASTEST (https://phastest.ca), a phage search tool web server (33).

### *E. coli* growth measurements

Overnight cultures of WT and Δ*cdt E. coli* strains in LB were subcultured 1:100 into LB, pooled human urine collected from healthy adult female volunteers, or RPMI 1640 (Gibco), and incubated at 37℃ with shaking. Growth was measured by CFU/ml and population doubling time (DT) was calculated as determined previously (45) from the measured CFUs at a time interval from 1h to 3h using the formula: DT=[Time duration(log2)]/[log(CFU at 3h)-log(CFU at 1h)].

To complement CFU kinetics and estimate bacterial growth *in situ*, the O:T^PCR^ method was used as described previously (34). Briefly, cultured bacteria were collected by centrifugation at 4℃ then immediately frozen on dry ice until further processing. Genomic DNA was extracted using the DNeasy Blood and Tissue kit (Qiagen) and quantitative PCR was performed for gene sequences near the origin (O) and terminus (T) of replication using the primer sequences provided in the table S1. This O:T^PCR^ method was also used to estimate the *E. coli* growth rates in mouse urine during experimentally induced UTI. For this application, urine from mice inoculated with wild-type HM56 and the HM56 Δ*cdt* mutant was collected at 6 and 24 h post inoculation, then processed as described for the *in vitro* samples.

### Gene expression measurements

HM56 was cultured overnight in LB, then sub-cultured in either pooled human urine or LB medium. Following incubation for 1 or 4 h at 37℃, bacteria were stabilized in RNAprotect (Qiagen), collected by centrifugation, and stored at -80℃ before RNA isolation. Total RNA was extracted from culture and infection samples using the RNeasy kit (Qiagen), reverse transcribed (iScript cDNA synthesis kit), and quantitative PCR was performed using the primer sequences SP9-12 (Table S1). Transcript levels of *cdtB* were normalized to the housekeeping gene, *gyrB*, and relative expression was calculated using the 2^-ΔΔCt^ method (46). Bacteria in LB cultures served as a reference condition for human urine.

### Mouse model of urinary tract infection

Female 6-7 week old CBA/J mice (Jackson Laboratory) were transuretherally inoculated with *ca*. 10^8^ CFU of *E. coli* as described previously (34). Urine from infected mice was routinely collected at 6 h post inoculation and daily beginning 24 post inoculation. The bladder and kidneys were harvested at 24 h or 7 days to quantify bacterial burdens.

To evaluate the relative fitness of HM56and the Δ*cdt* mutant derivative, mice were co-inoculated with a 1:1 mixture of both strains. Urine was collected after 24 h and then the bladder and kidneys were aseptically harvested and homogenized in sterile PBS. Bacterial burdens were determined from urine and tissue homogenates by plating samples on LB agar and LB agar supplemented with kanamycin, which selects for Δ*cdt* mutant bacteria. A competitive index (CI) was calculated using the formula: CI= (mutant CFU recovered/ wild-type CFU recovered) / (mutant CFU input/ wild-type CFU input). CI were log transformed and analyzed using one-sample t-test against the hypothetical neutral fitness value of 0 to assess statistical significance.

### Histopathology and Immunohistochemistry

Histology and Immunohistochemistry were performed by the University of Michigan Unit for Laboratory Animal Medicine Pathology Core (RRID:SCR_018823). Urinary bladder and kidneys were harvested at 1 or 7 days post inoculation from mice inoculated with either wild-type HM56, the Δ*cdt* mutant strains. Tissues were preserved in 10% neutral buffered formalin, then embedded in paraffin blocks. Bladders were bisected at the approximate level of the trigone and the kidneys were sectioned transversely on a rotary microtome at 4 µm thick. Slides were stained with hematoxylin & eosin for histological evaluation. Representative images were taken using an Olympus DP73 microscope camera and cellSens Entry 4.3 software. The severity and extent of inflammation in each section were scored in a blinded manner by a board-certified veterinary pathologist using semi-quantitative scoring scheme provided in Table S2 Briefly, the bladder and kidneys were each scored from 0-3 for the severity of cystitis or pyelonephritis, with zero signifying no significant lesions (NSL) and three indicating severe changes. Changes assessed included the type of inflammatory cell infiltrates, the relative numbers of inflammatory cells and their distribution throughout the tissue, and the relative degree of mucosal epithelial cell health. Infiltrates of polymorphonuclear cells (PMN) were weighted more heavily than lymphocytic / lymphoplasmacytic foci. Large rafts or aggregates of PMNs within the pelvis or bladder lumen were weighted more heavily than individualized cells, and samples with inflammatory cells disrupting or extending into the tissue parenchyma were weighted more heavily than those with only intraluminal cells. Mucosal changes included hyperplasia, vacuolation, apoptosis, and sloughing of epithelial cells. In the bladder, the extent of submucosal edema, inflammatory infiltrates, and peri-vascular cuffs were also included in the score weight. The complete histological scoring criteria, as well as representative images for a given score, are available in the supplementary materials (Table S2 and Fig. S5). For immunohistochemistry, the urinary bladder and a kidney from each mouse at 24 h were processed to paraffin, sectioned to glass slides, and stained with cleaved caspase-3 (Asp175) (Cell Signaling, #9661), 1:1000) antibody for evaluation of apoptosis. Reactivity score criteria are provided in Table S3.

## Acknowledgements

We acknowledge the members of our laboratory for their insightful feedback throughout this project. We also thank the histology technicians in the ULAM Pathology Core (RRID:SCR_018823) for their assistance in tissue processing and slide preparation.

This work was supported by funding from NIAID R01AI165582 (H.L.T.M, M.T.A., and M.M.P.).

